# Using citizen science data in integrated population models to inform conservation decision-making

**DOI:** 10.1101/293464

**Authors:** Orin J. Robinson, Viviana Ruiz-Gutierrez, Daniel Fink, Robert J. Meese, Marcel Holyoak, Evan G. Cooch

## Abstract

Analysis of animal population status and change are core elements of ecological research and critical for prioritizing conservation actions. Traditionally, count-based data from structured surveys have been the main source of information used to estimate trends and changes in populations. In the past decade, advances in integrated population models (IPMs) have allowed these data to be combined with other data sources (e.g., observations of marked individuals). IPMs have allowed researchers to determine the direction and magnitude of a species’ population trajectory and to test underlying mechanisms. For many species, life-history characteristics (e.g., low site-fidelity), low abundance and/or low detection probability make it difficult to collect sufficient data; thus, IPMs for these species are difficult to employ. We used count data from eBird to estimate population trends for one such species, the tricolored blackbird (*Agelaius tricolor*). We combined estimates of relative abundance with banding and nesting data. Our joint estimation of demographic rates allowed us to evaluate their individual contributions to the population growth rate. We suggest that investments in increasing reproductive success and recruitment are the most likely conservation strategies to increase the population. The extensive survey efforts of citizen scientists aided the employment of IPMs to inform conservation efforts.

**Data Accessibility:** Should the manuscript be accepted, the data supporting the results will be published along with the manuscript. The eBird data used for analyses in this manuscript may be downloaded from https://ebird.org/data/download.

## Introduction

The ability to generate information on the status and distribution of poorly known species has been a focus of recent developments in the fields of population ecology and conservation biology (Pacifici et al. 2012; Specht et al. 2017). Often, such species are rare, nomadic, have irruptive or unpredictable life histories, or possess other characteristics that make difficult the collection of counts and demographic information (Woinarski et al. 1992; Pedler et al. 2017). Count-based data across a species’ range are typically collected through survey programs that use a structured design (e.g., repeated counts; Royle 2004). This approach has worked well to characterize trends and distributions of populations at local (Keever et al. 2017) and continental scales (e.g. North American Breeding Bird Survey [BBS]; Sauer and Link 2011) for species that have predictable movements, or are faithful to specific sites at certain times of the year. However, most structured surveys are designed with a limited number of fixed sampling locations surveyed each year, which can lead to biased population trajectory/trend estimates for species with unpredictable intra- and/or inter-annual movements. That is, a high count in one year followed by a low count in the next may not be indicative of a decline, but rather that a significant number of individuals were observed at non-sampled sites during the second survey. Williams and Boyle (2018) showed that grasshopper sparrows (*Ammodramus savannarum*) changed sites, even within a breeding season, far more often than previously thought. This suggests that although an observer may be counting the same species at the same sites year after year, the likelihood of encountering the same individuals in successive years may be very low. As such, it may be difficult to infer population processes when using data collected in this manner.

Information on population vital rates is equally challenging to obtain relative to count data for nomadic or irruptive species, greatly limiting our ability to determine what factors influence population growth rates, or to define populations at all (Yoccoz et al. 2001). Mark-recapture studies are a widely used technique to estimate survival; however, species that do not return to the same breeding locations each year are unlikely to yield recapture rates that are high enough to allow for the robust estimation of survival and other vital rates (Marshall et al. 2004). For example, nomadic red crossbills (*Loxia curvirostra*) have low recapture probabilities and transient dynamics, which have been shown to underestimate survival (Alonso and Arizaga 2013). High rates of transience, such as those shown by American yellow warblers (*Setophaga petechia*), also lead to estimates of survival rates that are biased low (Cilimburg et al. 2002).

Integrated population models (IPMs) are increasingly being used in population ecology and have great potential as a wildlife management and conservation tool (Zipkin and Saunders 2018; Arnold et al. in press). IPMs allow researchers to jointly analyze data collected by different types of population studies (Schaub and Abadi 2011). Benefits of this joint analysis may be improved precision in the estimates of demographic parameters and population size, estimation of demographic parameters for which data are lacking, and simultaneous examination of the change in population size and the mechanisms underlying the estimated change (Kéry and Schaub 2012). While these models may accommodate many types of data, population counts or indices are required. These data allow the integration to occur, as the change in population over time is driven by the vital rates estimated by the other data sets in the model (Shaub and Abadi 2011). Population index data from national-scale surveys have been successfully used in IPMs to estimate large-scale population trends (e.g. Robinson et al. 2014; Ahrestani et al. 2017). However, these data come from rather strict sampling protocols conducted at fixed locations and applying them to nomadic or irruptive species may be more questionable.

Tricolored blackbirds (*Agelaius tricolor*) are nearly endemic to California, with no more than 1% of the population breeding in Washington, Oregon, Nevada, and the Mexican state of Baja California (Beedy et al. 2017). The species has declined rapidly, experiencing a 63% decline in its breeding abundance from 1935-1975 (Graves et al. 2013). This decline was concurrent with declines of freshwater wetlands and grasslands, their historical breeding habitat, although recent studies suggest that reproductive success may be as high or higher in alternative habitats such as Himalayan blackberry (*Rubus armeniacus*) and silage crops (Holyoak et al. 2014; Weintraub et al. 2016). The population has continued to decline since 1975, although the most recent California statewide survey (conducted in April 2017) suggests that the population may have increased slightly since 2014, or that non-breeding birds were uncounted previously and are now present in the breeding surveys (Beedy et al. 2017).

Tricolored blackbirds present monitoring challenges because of their unique life-history traits. They are a colonial breeding species, historically breeding in colonies as large as 200,000 individuals (Neff 1937). Carrasco et al. (2017) showed that colonial nesting herons and egrets strive to strike a balance between nest site fidelity and habitat preferences, suggesting that a colony may not form in a site that had previously been occupied. This has also been observed in tricolored blackbirds, where most colony sites are not used in consecutive years (Beedy and Hamilton 1997; Holyoak et al. 2014; Airola et al. 2017). This variable site occupancy can make it challenging to re-capture banded birds and/or to collect yearly population count data (Marshall et al 2004). As most surveys are conducted at fixed locations, it may be difficult to interpret year-to-year counts of a species that is not likely to be in the same count location for each subsequent sampling occasion. Early efforts for counting tricolored blackbirds have summed the number of birds across all breeding sites, however this metric was shown to be highly dependent upon the number of colony sites sampled and is not recommended (Graves et al. 2013). The current method is a coordinated range-wide survey conducted triennially (e.g. Meese 2014) and augmented by a sampling of the statewide population in years where the larger survey does not occur. However, these efforts are focused on previously occupied sites and may miss colonies that are established in new locations.

Given the uncertainties associated with tricolored blackbird count and index data (e.g., BBS), we used data from the eBird citizen-science project (Sullivan et al. 2014) as the population index data in an IPM. These data provide comprehensive spatial coverage and are independent of species occurrence as they may be collected anywhere in a species’ range. As such, they may provide the best available representation of the population trajectory over the last 10 years of this highly unpredictable species. We integrated eBird data for tricolored blackbirds from the pre-breeding period with band re-capture data from 18 sites and fecundity data from 10 sites in an IPM to determine the population trajectory and to identify life-history stages that show the most potential for increasing populations. This study provides the first use of less structured, citizen-science data in an IPM, and emphasizes the potential for these types of rapidly growing databases to become powerful tools in the fields of applied ecology and conservation biology.

## Methods

### Data

Tricolored blackbirds were banded at 18 sites throughout central California using United States Geological Survey (USGS) metal bands from 2007-2016. Trapping occurred during the breeding season (April – July) using walk-in traps, typically baited with cracked corn infrequently supplemented with mealworms. Recaptures were recorded each year during the breeding season. Each breeding season was considered a separate recapture occasion and data for within-season recaptures were not included in our estimates of annual survival. We used data from 64,129 tricolored blackbirds (49,668 females and 14,461 males) banded and 1,460 recaptured (1,336 females and 124 males) as adults in our analysis.

As raw fecundity data were unavailable, we simulated the fecundity data based on the reported summary statistics for fledgling surveys from 1992-2016. This was conducted in such a way to capture the substantial year-to-year variation expected in the underlying data. As between 60 and 100 nests (*n*) were surveyed in each year (*i*) per site (*s*), we allowed the number of nests sampled in a year to be a random draw from a uniform distribution with minimum 60 and maximum 100. The number of sampled sites varied among years between two and 10, thus we allowed the number of sites to be drawn from a uniform distribution with a minimum of two and maximum of 10. We then calculated the long-term mean and standard deviation of the annual assumed fledglings (defined as 5-9 day old chicks) per nest (*h*) from 1992-2016. We calculated the proportional difference for the years 2007-2016 relative to the long-term average of fledglings produced per nest in order to modify the estimates for these years.

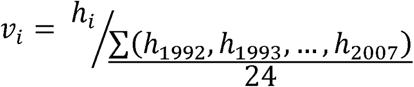

Thus, *v* is a measure of how far the number of fledglings per nest is from the long-term average, scaled to one. If *h* for a given year was exactly the long-term average, it was multiplied by one. If it was above or below the average, it was multiplied proportionally to how far above or below the average it was. We calculated the number of fledglings (*J*) counted in year *i* as

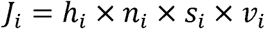

This was simulated 1000 times and the mean values for *J* and *n* were used as data for the IPM.

We used relative abundance calculated using counts of tricolored blackbirds recorded in eBird as the count data for the IPM. We collected eBird records for tricolored blackbirds across all of California for the years 2007-2016 and during the early breeding period (March 23-April 25; Additional details about eBird data selection and preparation are in Appendix 1). We only kept checklists that recorded an abundance or absence for tricolored blackbirds. We removed checklists simply noting occurrence without an estimated abundance. After implementing these constraints, 113,501 checklists remained. We separately filtered the data by year according to Robinson et al. (2017) in order to balance the data set temporally and spatially, and to correct for class imbalance. As recent years contain far more checklists than older ones, we restricted the number of samples per year to control for the increase in eBird participation. We set 2007 as our reference year, fixing the sample size of the spatially balanced data for each year to equal that of 2007. As this removed some checklists from the data for the years after 2007, we resampled across all of the checklists 1000 times, creating 1000 unique eBird data sets, filtered from the original 113,501 checklists that remained.

Following Fink et al. (2018), we used each of these data sets in a two-step zero-inflated generalized additive model (ZI-GAM; Appendix 2) to estimate the relative abundance of tricolored blackbirds for each year of the study. Parameters were low-order piecewise polynomial basis functions with limited knots to restrict the flexibility of the spline components, thereby controlling the bias-variance tradeoff and improving the stability of the ZI-GAM. The first step is a binary response (*Ψ*) of detection (1) or non-detection (0) and had the form

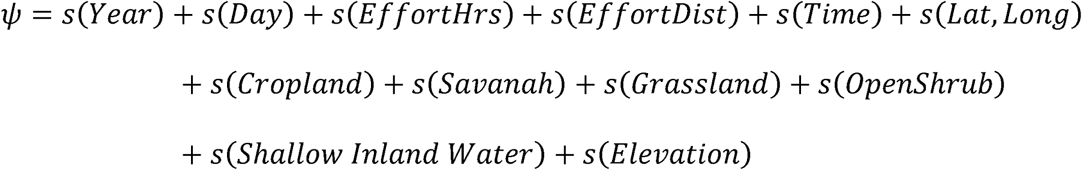

The second step of the ZI-GAM is a Poisson response of the estimated abundance of tricolored blackbirds on an eBird checklist, given that a tricolored blackbird was detected in the first step, and took the form

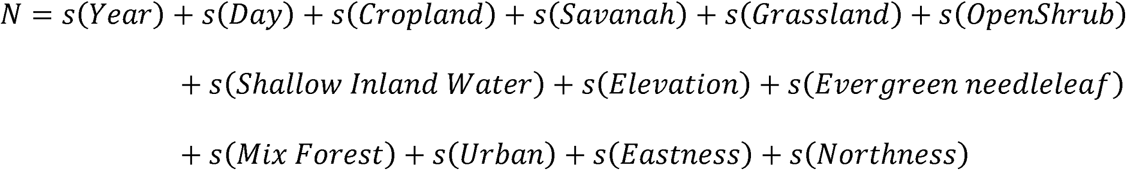

Covariates for time spent searching (*EffortHrs*), distance traveled while searching (*EffortDist*), and time of day (*Time*) in the detection portion; and year (*Year*), and day of the year (*Day*) in both portions were included to account for variation in detectability and variation in availability for detection. Coarse-scale spatial variation was captured by a smoothed two-dimensional function of latitude and longitude (*Lat, Long*) and fine-scale spatial variation was captured by the MODIS land cover classes. To account for variation in the spatiotemporal balancing step described above, we fit the ZI-GAM to each of the 1000 replicate data sets and used the mean result in the analysis. These models were fit using the mgvc (Wood 2011) package in R (R Core Team 2016).

### Integrated population model

We analyzed the demographic data described above using a sequential method of the ZI-GAM followed by an integrated population model (Besbeas 2002; Schaub and Abadi 2011). We based our IPM on a pre-breeding, 2×2 stage-based projection model and three separate sub-models to link adult survival for both males (*ϕ*_*am*_) and females (*ϕ*_*af*_), juvenile survival (*ϕ*_*j*_), fecundity (*F*) and relative abundance (*N*_*tot*_). We modeled relative abundance (*N*) for both age classes (non-adults [1] and adults [2]) at each year (*t*) so that

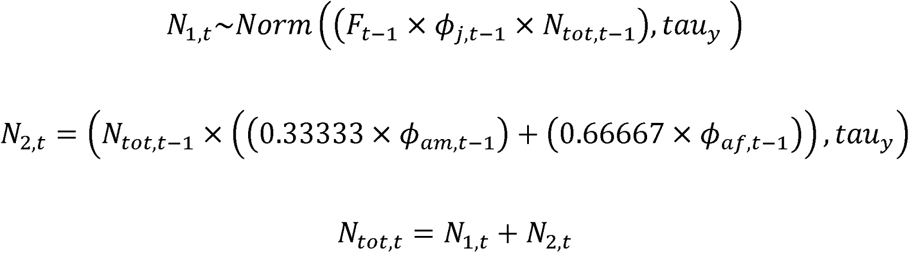

where *tau* is the estimated error in the count data given an uninformative uniform prior (min = 0 and max = 10,000), *F* is estimated from the fecundity data

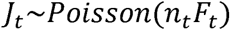

and *ϕ* is estimated for adults of each sex independently using the multinomial likelihood of the CJS model (Kéry and Schaub 2012). As tricolored blackbird colonies have a sex ratio skewed towards females, we were concerned that if true differences in survival exist, assuming equal survival probabilities between the sexes may bias the results (Gerber and White 2013). Male adult survival was modified by 1/3 and female adult survival by 2/3 as there is roughly one male for every two females in a colony, however there may be between one and four females per male (Beedy et al. 2017). As we only used banding data for adults, *ϕ*_*j*_ was a latent parameter of the model and we assumed that juvenile survival did not differ between males and females. We did not include immigration as our abundance data from eBird spanned the entire state of California, which includes roughly 99% of the breeding population (Beedy et al. 2017). We estimated the growth rate (*r*) of the population during the 10 years of the study from the posterior estimates of relative abundance. We then compared the estimates of *ϕ*_*af*_, *ϕ*_*am*_, and *F* with *r* to determine the effect of each vital rate on population growth via correlation analysis. Posterior distributions of the demographic parameters and relative abundances were estimated from 50,000 iterations after a burn-in of 5,000 samples. Convergence was considered successful if R hat values were less than 1.1 for all parameters (Gelman and Rubin 1992). The IPM was fit using the package r2jags (Su and Yajima 2015) in R (R Core Team 2016).

## Results

All credible intervals (CI) reported are the 95% highest posterior density intervals. The relative abundance in the analysis of eBird data alone estimated a decline of 52%, with the highest relative abundance within the timeframe of the study occurring in 2007 and the lowest occurring in 2013 (Figure 1). The IPM analysis estimated a mean decline of 34% (CI = 7.5% growth to 71% decline), with the highest relative abundance occurring in 2008 and the lowest in 2014 (Figure 1). The growth rate of the population was negative (mean −0.06, CI −0.14 – 0.016) although the CI interval slightly overlapped zero. This is highly suggestive of a decline, as 94% of the IPM iterations resulted in an estimated growth rate below zero (Figure 3). Females had a higher overall mean apparent annual survival than males (Table 1) although the credible intervals for most year-interval estimates overlapped between males and females (Figure 2). Yearly estimates for juvenile survival were uninformative, having credible intervals for some years that spanned the entirety of the allowable parameter space (i.e. 0-1). Therefore, we calculated a long-term mean juvenile survival across years (mean 0.21, CI 0.0007 – 0.49). Mean yearly fecundity spanned values from 0.46 fledglings per nest in the least productive year to 1.27 in the highest (Table 1).

**Figure 1.**
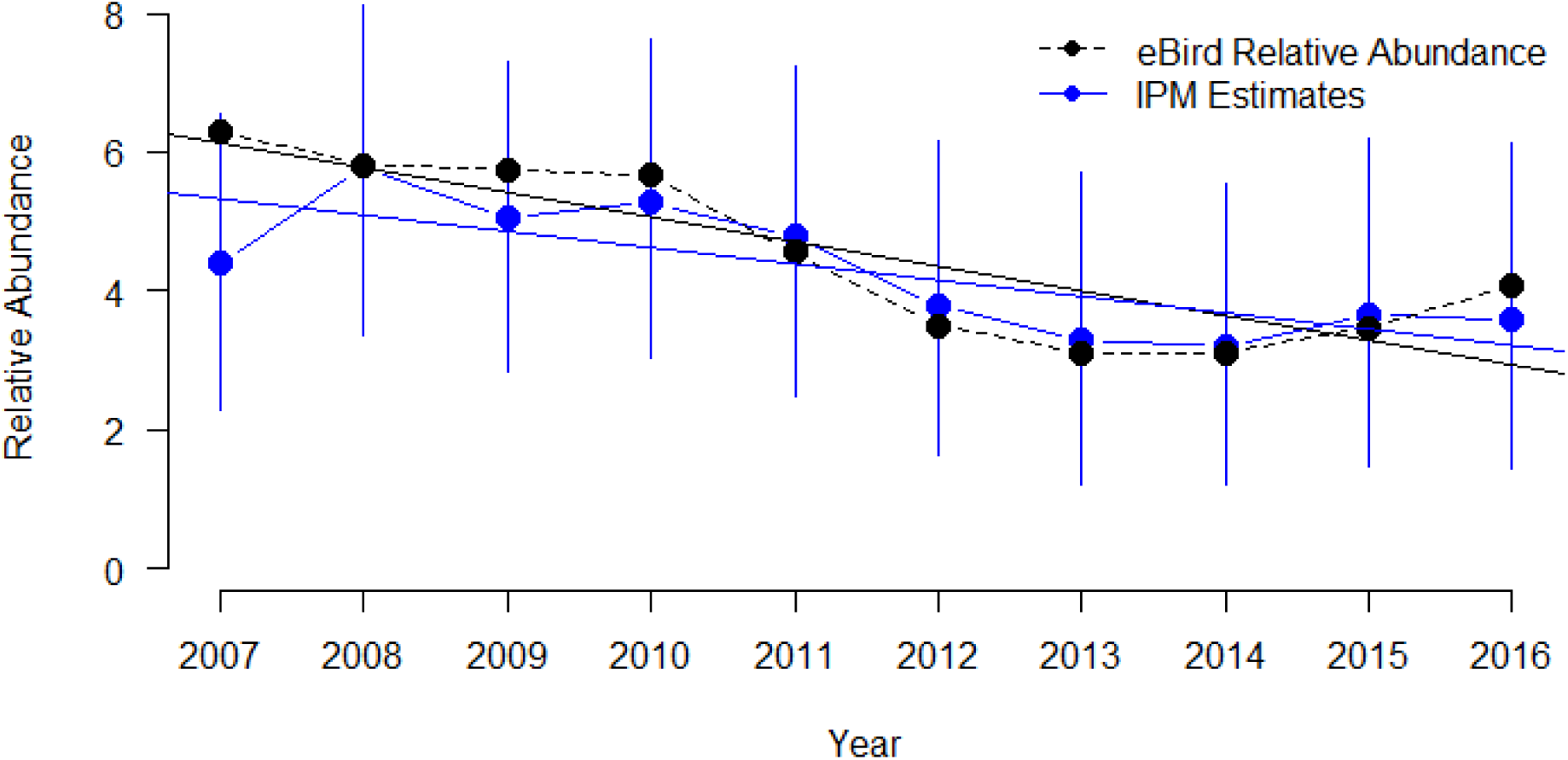
Estimates of relative abundance from eBird (black) and the integrated population model (blue) with 95% highest posterior density intervals. The straight lines are predicted values from a GLM on each time series.

**Figure 2.**
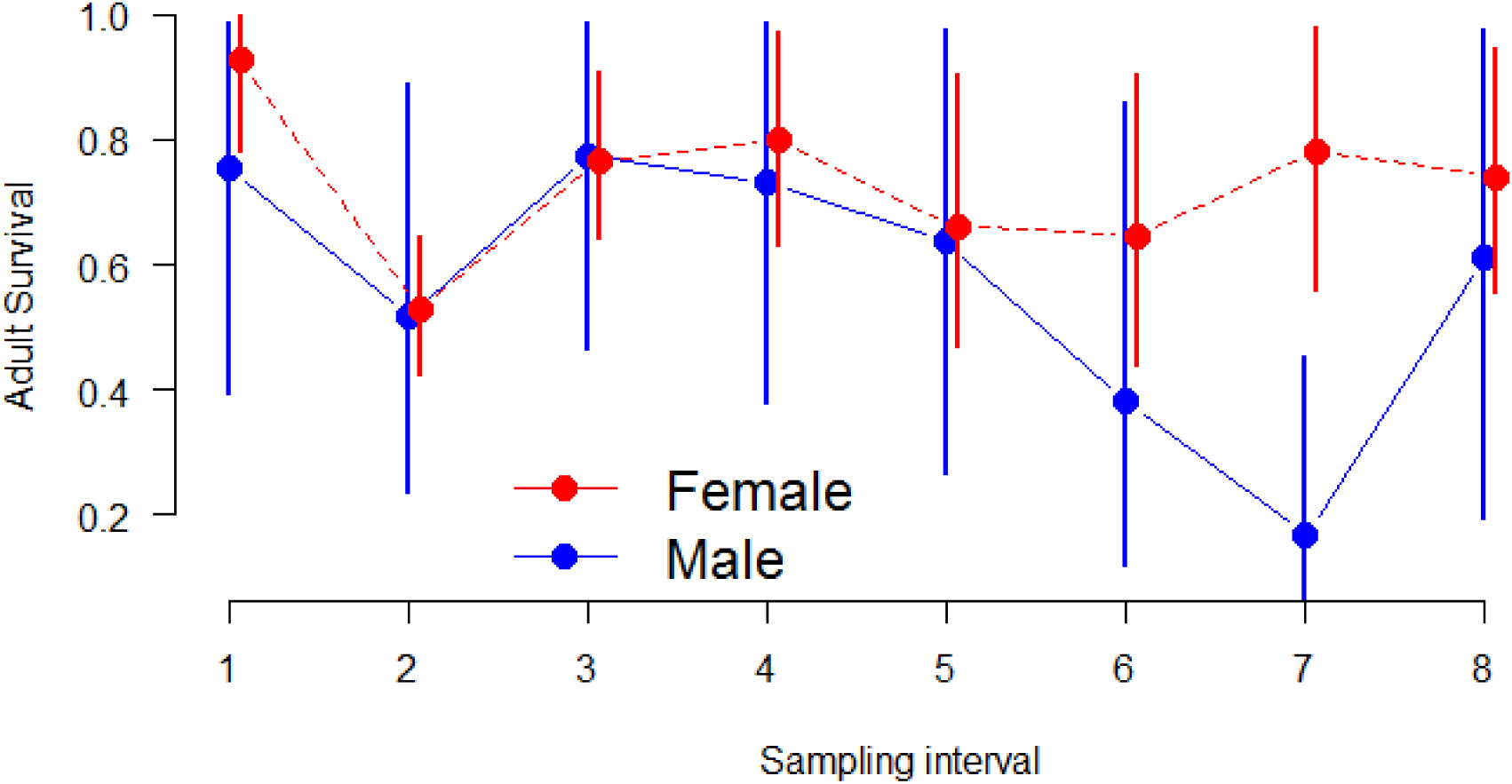
Adult survival estimates and 95% highest posterior density intervals for female (red) and male (blue) tricolored blackbirds for each sampling interval (i.e., the probability an adult survived during the time-period between each sampling occasion). Interval 1 is the period between banding field seasons of 2007-2008, interval 2 is between 2008-2009, etc. The last sampling interval (9) has been removed because survival and recapture probabilities are confounded for the last interval and cannot be estimated separately.

**Figure 3.**
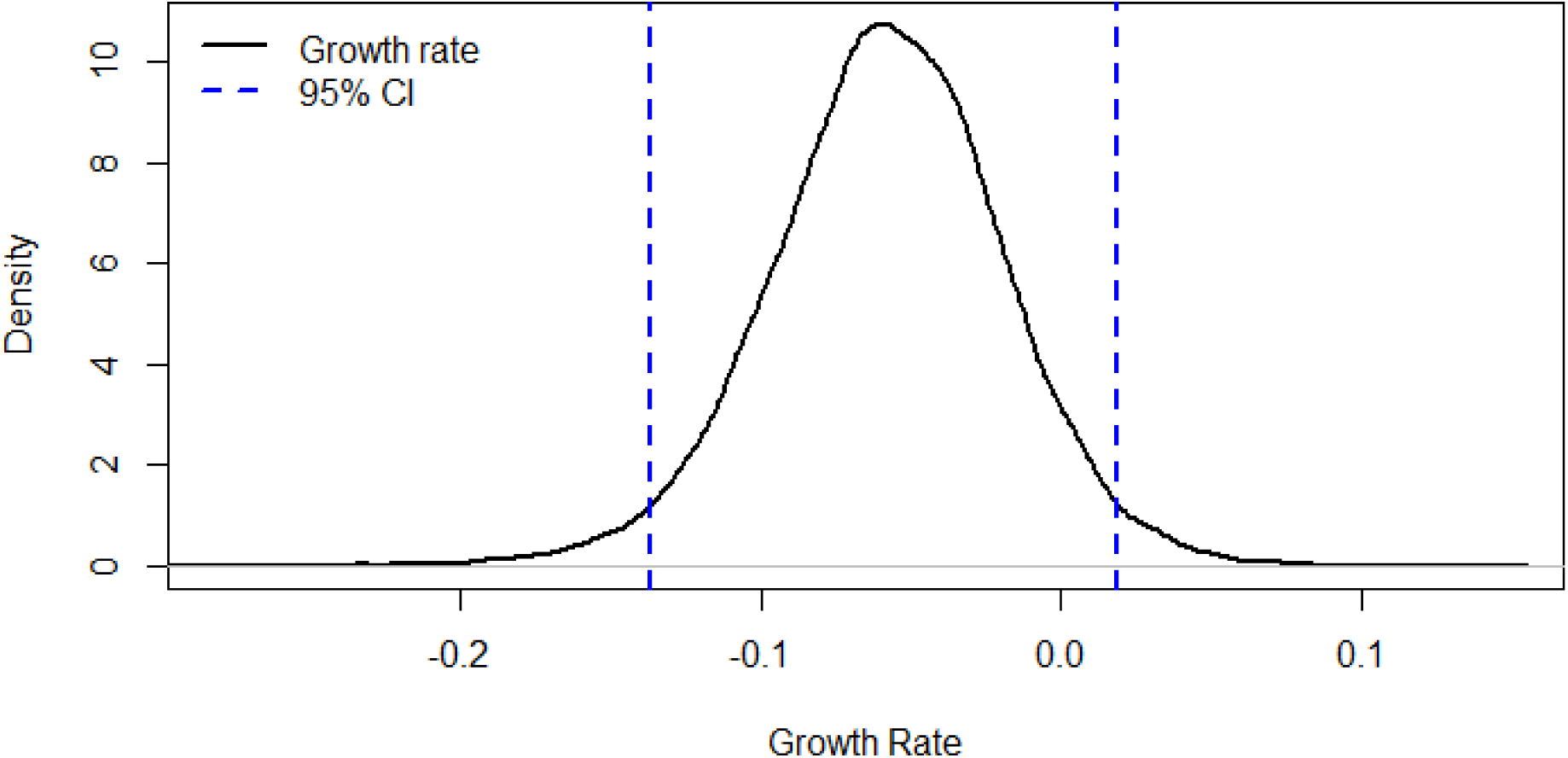
Posterior distribution of the growth rate estimated by the integrated population model. Vertical dashed blue lines represent the 95% highest posterior density intervals.

**Table 1.**
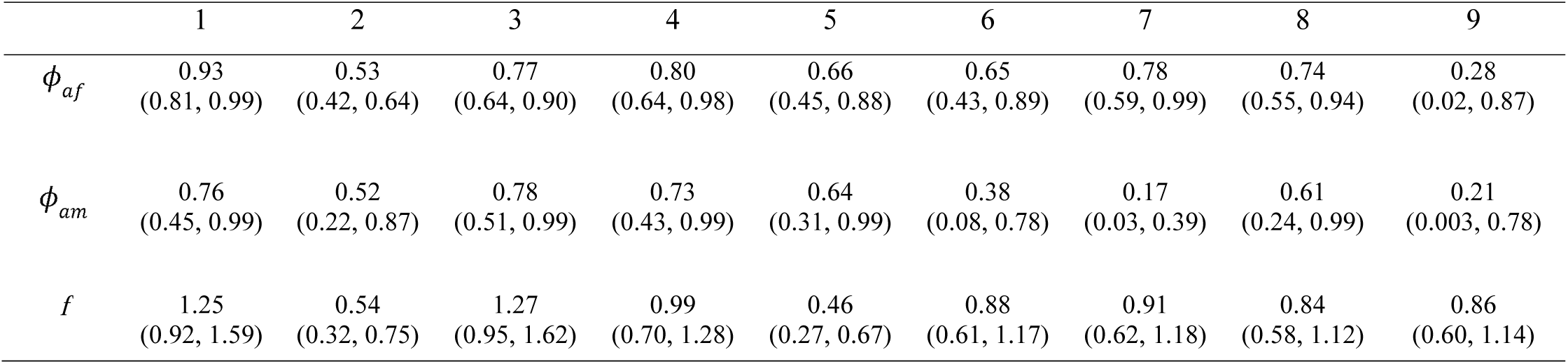
Estimates of annual adult female survival (*ϕaf*), adult male survival (*ϕ*_*am*_), and fecundity (*f*) for each time interval during the study with the highest posterior density interval in parentheses. Interval 1 is the period between banding field seasons of 2007-2008, interval 2 is between 2008-2009, etc.

|  | 1 | 2 | 3 | 4 | 5 | 6 | 7 | 8 | 9 |
| --- | --- | --- | --- | --- | --- | --- | --- | --- | --- |
| $\phi_{af}$ | 0.93<br>(0.81, 0.99) | 0.53<br>(0.42, 0.64) | 0.77<br>(0.64, 0.90) | 0.80<br>(0.64, 0.98) | 0.66<br>(0.45, 0.88) | 0.65<br>(0.43, 0.89) | 0.78<br>(0.59, 0.99) | 0.74<br>(0.55, 0.94) | 0.28<br>(0.02, 0.87) |
| $\phi_{am}$ | 0.76<br>(0.45, 0.99) | 0.52<br>(0.22, 0.87) | 0.78<br>(0.51, 0.99) | 0.73<br>(0.43, 0.99) | 0.64<br>(0.31, 0.99) | 0.38<br>(0.08, 0.78) | 0.17<br>(0.03, 0.39) | 0.61<br>(0.24, 0.99) | 0.21<br>(0.003, 0.78) |
| $f$ | 1.25<br>(0.92, 1.59) | 0.54<br>(0.32, 0.75) | 1.27<br>(0.95, 1.62) | 0.99<br>(0.70, 1.28) | 0.46<br>(0.27, 0.67) | 0.88<br>(0.61, 1.17) | 0.91<br>(0.62, 1.18) | 0.84<br>(0.58, 1.12) | 0.86<br>(0.60, 1.14) |

The years of highest relative abundance followed the intervals where fecundity and female survival were also at their highest. Adult female survival and fecundity were positively correlated with population growth rate (ρ = 0.24, CI −0.50, 0.77 and ρ = 0.33, CI −0.45, 0.76 respectively) and the mean posterior probability that each correlation coefficient is positive was 0.70 and 0.69. Adult male survival was slightly and negatively correlated with growth rate (ρ = - 0.06, CI −0.61, 0.67), and its probability of being positively correlated with the growth rate was 0.55.

## Discussion

As an array of factors continues to threaten wildlife populations, it is imperative to understand the drivers of population dynamics. IPMs provide a framework by which population growth and the processes driving it can be simultaneously explored (Kéry and Schaub 2012; Duarte et al 2017). One requirement in IPMs is a reliance upon abundance data of the species of concern, which may be difficult to acquire for species with complex life histories. We have demonstrated an approach to sequentially model abundance data from eBird, and incorporate this into an IPM to analyze population trend, band recapture, and fecundity data for a species of conservation concern. We were able to explore the population trend of tricolored blackbirds and its response to fecundity and adult survival and to estimate juvenile survival, a parameter for which we have no data. This analysis would not have been possible without making use of eBird data, as the more statistically robust surveys have only occurred for the last three triennial Statewide Surveys.

Our estimates of relative abundance from the IPM closely follow the relative abundance estimates from eBird data, although the models disagree on the year of highest relative abundance. In this analysis, eBird data cannot give a measure of absolute abundance, so our results can only be interpreted as an estimate of a population trend, and not population size at each year. Our analysis indicates that the population of tricolored blackbirds has likely declined between 2007 and 2016, as 94% of the IPM iterations resulted in a growth rate below zero. We also estimated that the magnitude of that decline is 34%. Visual inspection of our estimated population trajectory (Figure 1) appears to show that the population may have somewhat stabilized in the last few years of the study, a result in accord with the results of the most recent Statewide Survey of the species (Meese 2017). There are several possible interpretations of this result. First, although the population may have appeared to cease to decline from 2013 onwards, it may have been because more birds attempted to breed from 2013 onwards than immediately prior to this; that is, proportionately more birds were present as nonbreeding individuals prior to 2013 and attempted to breed from 2013 onwards. Second, recruitment may have increased in more recent years, perhaps because a large number of breeding birds being protected in grain fields due to dairy farmers being compensated to delay crop harvest to allow birds to fledge (Aurthur 2015; Meese 2017), or because adult or juvenile survival increased in more recent years. Neither the reproductive success nor survival ideas are supported by the results of our IPM analyses for these specific years, although the overall result suggests that fecundity and adult female survival have large effects on the population growth rate. Our eBird data may be missing some counts from colonies on private land. Hence, it is possible that they do not capture results for a portion of the population, such as colonies breeding in private grain fields. We also caution that a small but stable population does not necessarily mean a healthy one (Lacy 2000; Robles and Ciudad 2012).

It is common for population models to model only females (Caswell 2001), or to assume both males and females have equal survival probabilities (e.g. Robinson et al. 2016; Saunders et al. in press). As we were concerned that the skewed sex ratio in tricolored blackbird colonies may bias our results, we estimated female and male adult survival separately within the IPM. We also varied the proportion of males and females in the model (results not shown) and determined that the ratio assumed did not make a difference to the population trend for tricolored blackbirds or the survival estimates until the ratios were extremely skewed (≤ 10% of either sex). We found that mean adult female survival is generally higher than male survival, although the credible intervals overlapped for most estimates. Our results also suggest higher average annual adult survival estimates than previously unpublished analyses (0.6; Beedy et al. 2017).

The results of our correlation analysis suggest that conservation action on either female survival or fecundity may achieve the greatest benefit to the population as they had similar contributions to population growth rate. However, adult female survival is higher than previous estimates have suggested (Beedy et al. 2017) and similar to adult survival estimates of red-winged blackbirds (*Agelaius phoeniceus*; Frankhauser 1967) and yellow-headed blackbirds (*Xanthocephalus xanthocephalus*; Bray et al. 1979). As this vital rate is already comparatively high, there may not be room to improve it via conservation measures. Therefore, conservation practitioners may consider actions that improve fecundity, as the majority of years for which we estimated fecundity were below one chick per nest. Conservation action to protect nest sites and improve fecundity are already underway in California, where farmers are being compensated to offset lost crop value from delaying harvest of the grain crop in which tricolored blackbirds are nesting (Arthur 2015). As conservation is dynamic, more research will be required to assess the effectiveness of these efforts, and to adjust conservation measures as needed (Reynolds et al. 2017).

The use of IPMs for wildlife management and conservation is growing (Arnold et al. in press; Zipkin and Saunders 2018), as is the number of participants in citizen science data collection (Theobald et al. 2015). Citizen science data have been used to enrich existing data sets on species occurrence and abundance (e.g. Pacifici et al. 2017) and the growth of citizen science programs may facilitate their integration with other data types as well. For species, like tricolored blackbirds, that may be challenging to monitor through more traditional schemes, data in eBird can be used to estimate large-scale population trends and life-history parameters. Observations collected in eBird helped to alleviate the issues associated with counting species with low site fidelity as it is an open platform and, thus, data may be collected at any site within the species’ range. Because surveys can be conducted at any location, eBird can cover much larger spatial and temporal extents and be more flexible surveying areas compared to traditional surveys. Data integration and rapidly growing citizen science databases are poised to become powerful tools in conservation science (Zipkin and Saunders 2018). Here, we have taken a first step to combine citizen science data with recapture and fecundity data in an IPM to help inform more effective conservation actions for tricolored blackbirds. Our research and recommendations for conservation strategies were a direct result of the joint analysis that would not have been possible without making use of citizen science data.

## Appendix 1 eBird data selection and preparation

The tricolored blackbird abundance information came from the citizen-science project eBird (Sullivan et al., 2014). eBird is a global bird monitoring project that collects observations made throughout the year. Participants follow a checklist protocol, in which time, location, search effort and counts of birds are all reported in a standardized manner. By asking participants to indicate when they have contributed ‘complete checklists’ of all the species they detect and identify on a search, we can assume failure to report a species conveys information about detection probability and presumed absence of the species. We used the inferred counts of zero birds, together with information on observers’ effort, to account for sources of variation associated with the detection process.

There are a number of specific data fields that participants fill in when they submit observations that aid in how data is filtered. First, participants collected observations via checklists that contain fields for location, time (day, month, year, time of day), distance travelled, duration, number of observers, and counts of species encountered. The checklists also pass through automated filters that flag anything that may be unusual, such as an observation of a species that is rare at a specific location and/or time of year, or an abnormally high count of a species (Sullivan et al. 2014). A network of more than 500 experts around the world manage the checklists submitted by birders, working to verify any checklist flagged by the automated filters.

Data for the ZI-GAM was from a subset of complete checklists collected under the ‘travelling count’ and ‘stationary count’ protocols. These data were generated from checklists with transect distances ≤ 8.1 km, start times during daylight hours between 0500 and 2000 local time, and total search times ≤ 6 h. Any checklist with an ‘X’ in the count field was removed.

## Appendix 2 Zero-Inflated Generalized Additive Model and ZI-GAM code

The ZI-GAM is essentially two models that ask the questions 1) “Is there a tricolored blackbird detected?” and 2) “If so, how many?”

In the detection portion, variables associated with detection in eBird are used as predictors. Those are effort in terms of hours spent searching and distance travelled, time variables (year, day, and time of day), elevation, slope direction (Eastness and Northness) and MODIS land-cover variables measured every two weeks.

In the count portion, day, year, elevation, slope direction and land-cover variables are included as predictors.

The land-cover variables are those that are associated with tricolored blackbirds (taken from Robinson et al. 2018) and used in one or both portions of the ZI-GAM: Croplands, Savanah, Grasslands, Open Shrublands, Shallow Inland Water, Evergreen Needleleaf (negative but strong association), Mixed Forest, Urban and Built-Up.

Example code for ZI-GAM

~~~
d.gam <-gam(list(
### This is the count portion of the GAM ###
Agelaius_tricolor ~
s(DAY, m=1, k = 10) + s(YEAR, m=1, k=4) + s(Croplands, m=1, k=5) + s(Savanah, m=1, k=5) +
s(ELEV, m=1, k=5)+ s(Grasslands, m=1, k=5)+
s(Open Shrublands, m=1, k=5)+s(Shallow inland water, m=1, k=5)+
s(EASTNESS, m=1, k=5)+ s(NORTHNESS, m=1, k=5) + s(Evergreen needleleaf, m=1, k=5) +
s(Mixed forest, m=1, k=5) + s(Urban & built up, m=1, k=5),
~
### This is the detection portion of the GAM ###
s(DAY, m=1, k = 10) +
s(EFFORT_HRS, m=1,k=k.ttt) + s(YEAR, m=1,k=4) +
s(EFFORT_DISTANCE_KM, m=1,k=k.ttt) +
s(TIME,m=1,k=k.ttt) + s(LATITUDE,LONGITUDE)+
s(Croplands, m=1, k=5) + s(Savanah, m=1, k=5) +
s(ELEV, m=1, k=5)+ s(Grasslands, m=1, k=5)+
s(Open Shrublands, m=1, k=5)+s(Shallow inland water, m=1, k=5)),
family = ziplss,
gamma = 1.4,
data =Spatially filtered eBird data set)
~~~

